# Comprehensive analysis of the host-virus interactome of SARS-CoV-2

**DOI:** 10.1101/2020.12.31.424961

**Authors:** Zhen Chen, Chao Wang, Xu Feng, Litong Nie, Mengfan Tang, Huimin Zhang, Yun Xiong, Samuel K. Swisher, Mrinal Srivastava, Junjie Chen

## Abstract

Host-virus protein-protein interaction is the key component of the severe acute respiratory syndrome coronavirus 2 (SARS-CoV-2) lifecycle. We conducted a comprehensive interactome study between the virus and host cells using tandem affinity purification and proximity labeling strategies and identified 437 human proteins as the high-confidence interacting proteins. Functional characterization and further validation of these interactions elucidated how distinct SARS-CoV-2 viral proteins participate in its lifecycle, and discovered potential drug targets to the treatment of COVID-19. The interactomes of two key SARS-CoV-2 encoded viral proteins, NSP1 and N protein, were compared with the interactomes of their counterparts in other human coronaviruses. These comparisons not only revealed common host pathways these viruses manipulate for their survival, but also showed divergent protein-protein interactions that may explain differences in disease pathology. This comprehensive interactome of coronavirus disease-2019 provides valuable resources for understanding and treating this disease.

## INTRODUCTION

The ongoing global pandemic of coronavirus disease-2019 (COVID-19) was first reported in December 2019. Severe acute respiratory syndrome coronavirus 2 (SARS-CoV-2) was identified as the virus causing COVID-19^1^. SARS-CoV-2 is a highly transmissible and pathogenic coronavirus. It spreads easily through the air when people are physically near each other. As of 18 December 2020, more than 75 million cases had been confirmed, resulting in more than 1.6 million deaths.

SARS-CoV-2 is a novel beta coronavirus with a genome composed of approximately 30 kb of positive-strand RNA. It shares 79% genomic sequence identity with SARS-CoV-1, which caused the SARS epidemic in 2003^2^. The SARS-CoV-2 genome contains 14 open reading frames (ORFs), including one large ORF that encodes two large polyproteins (ORF1a and ORF1ab) and 13 small ORFs that encode viral structural proteins and other polypeptides. The polyproteins from the large ORF can be further cleaved into 16 non-structure proteins (NSP1 to NSP16)^2^.

Researchers have tried several strategies to develop drugs and vaccines to treat SARS-CoV-2 and COVID-19. Viruses usually encode very limited viral genes and proteins and thus need to recruit many host proteins to complete the viral lifecycle. Therefore, identifying proteinprotein interactions between viral proteins and their host cellular cofactors is an important and efficient way to understand the virus and uncover potential drug targets. Such host-virus interactome analysis has been reported recently^3^.

There are two well-developed strategies to study the protein-protein interactome, affinity purification (AP) and a proximity labeling-based strategy, followed by mass spectrometry (MS) analysis^4^. AP-MS is a widely used and highly reproducible method that allows identification of physiologically relevant interaction proteins. However, this method may miss weak or transient binding proteins during the pulldown process, even when it is performed in a mild buffer. In addition, AP-MS may not capture poorly soluble protein partners, such as membrane-associated proteins, which are critically important for the viral lifecycle. The proximity labeling-based strategy solves these problems by labeling nearby proteins before harvesting cells. This method is based on an enzyme-substrate reaction with an effective labeling radius of about 10 nm. The drawback of this method is that it may fail to label the binding proteins located outside of this labeling range. Therefore, combining these two strategies should allow us to generate a comprehensive host-virus interactome network that is important for the SARS-Cov-2 lifecycle.

In the current study, we applied these two strategies, i.e., tandem affinity purification (TAP) with the SFB (S-protein, FLAG epitope, and streptavidin-binding peptide) tag and proximity labeling with a second-generation biotin ligase, BioID2, for a comprehensive analysis of the host-virus protein-protein interaction network. With an interactome analysis of 29 SARS-CoV-2 cDNAs, we uncovered key human proteins involved in the SARS-CoV-2 lifecycle of infection, replication, and budding. This interactome dataset not only confirmed some previously reported host-virus interactions but also uncovered numerous new interacting proteins that may be critical for the SARS-Cov-2 lifecycle. This dataset will benefit further investigations of the mechanisms underlying viral infection and lifecycle and potential new drug targets for the treatment of COVID-19. Moreover, we compared the interactomes of two critical viral genes, NSP1 and N protein, among different human coronaviruses. These analyses will help us to fight SARS-CoV-2 and future pandemics caused by new coronaviruses.

## RESULTS

### Overview of the Interactome Analysis between Host and SARS-CoV-2

To comprehensively illustrate the host and SARS-CoV-2 protein-protein interaction network, we performed an interactome study using two different strategies, TAP with the SFB tag and proximity labeling with the BioID2 tag. Genome annotation revealed 29 gene products from SARS-CoV-2, including 16 NSPs, 4 structure proteins, and 9 accessary factors (**Figure 1A**). These viral gene products were fused with the SFB or BioID2 tag and stably expressed in HEK293T cells, except NSP1, which was done with transient expression. Stable viral gene expression in cells was verified by immunoblotting (**Figures S1A and S1B**). Labeling reactions of fused BioID2-viral genes were confirmed using the streptavidin antibody (**Figure S1C**). Three biological replicates of the interactome experiments were conducted for each tag and each fused viral gene, along with controls (vector with tag only or fused with GFP sequence) following the workflow presented in **Figure 1B**. After analysis by Q Exactive HF MS, the data were searched against a database integrated with all human genes, viral genes, GFP, and the tag sequences (**Tables S1**). All experiments are summarized in **Figure 1C**. The Pearson correlation coefficient among three independent biological replicates of the SFB-TAP results and BioID2 labeling experiments was calculated (**Figure 1D**).

**Figure 1.**
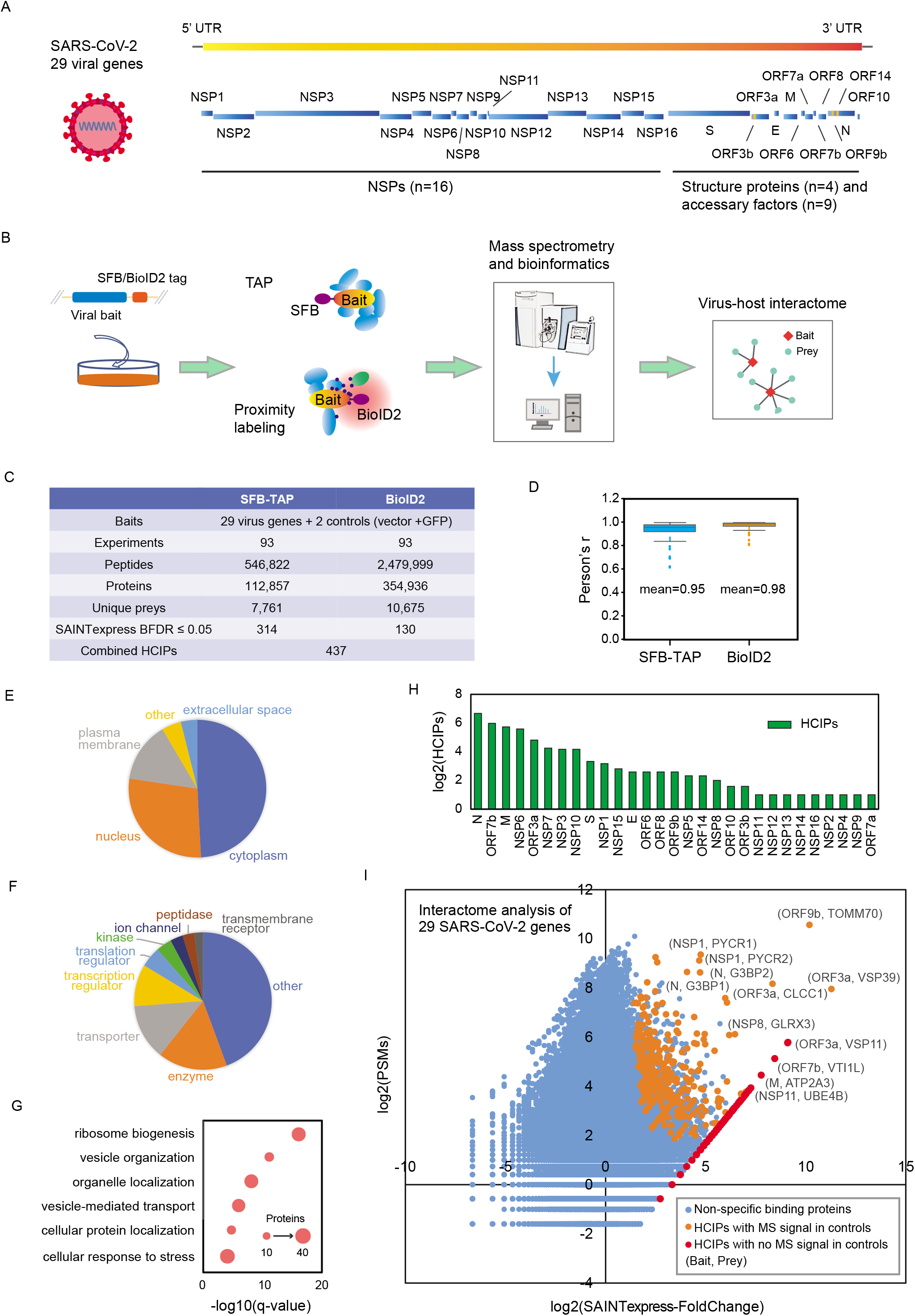
Summary of the SFB-TAP and BioID2 interactome experiments. (A) SARS-CoV-2 genome annotation, predicting 29 virus gene products. The 16 non-structure proteins (NSPs) are cleaved products of the large polyprotein open reading frame (ORF)1ab or ORF1a. These polyproteins are cleaved into small function fragments or NSPs after translation. (B) Workflow for the comprehensive virus-host interactome analysis. Two different labeling strategies, SFB-TAP and BioID2 labeling, were applied in the study. Samples were analyzed by Q Exactive HF mass spectrometry (MS). (C) Summary of the datasets obtained from SFB-TAP and BioID2 results, including the number of high-confidence interacting proteins (HCIPs). BDFR, Bayesian false discovery rate. (D) Pearson correlation coefficient among three independent biological replicates of the SFB-TAP results and the BioID2 labeling experiments. (E-G) GO analysis. GO enrichment was performed using Ingenuity Pathway Analysis. Protein localization (E), molecular function (F), and biological function (G) are plotted in a single panel. (H) HCIPs identified in the purification of each SARS-CoV-2 gene. (I) Correlation between peptide-spectrum matches (PSMs) of identified proteins and their fold change calculated by SAINTexpress.

The identified proteins were filtered using Significance Analysis of INTeractome (SAINTexpress)^5^. The SAINTexpress scores were averaged among the three biological replicates to calculate a Bayesian false discovery rate, and preys with a Bayesian false discovery rate of 0.05 or less were considered high-confidence interacting proteins (HCIPs). In total, we obtained 314 HCIPs from SFB-TAP and 130 HCIPs from BioID2 experiments (**Table S2**). We combined these two datasets and generated a list of HCIPs, which included 437 pairs of virus-host interacting proteins (**Figure 1C**). GO analysis of these 437 viral interacting proteins showed that the preys were localized in various subcellular locations (**Figure 1E**). Many of the preys localized on various plasma membrane structures, such as the endoplasmic reticulum, Golgi membrane, or cell membrane. Protein function analysis also showed that many of the preys had roles in transporter, ion channel, and peptidase processes (**Figure 1F**). The biological processes enriched included vesicle organization, organelle localization, and vesicle-mediated transport (**Figure 1G**). These preys may assist SARS-CoV-2 infection, replication, and budding. Another group of enriched proteins is involved in ribosome function, indicating that these preys may be exploited by viral proteins to suppress host cell translation and facilitate viral gene expression.

The number of HCIPs for each SARS-CoV-2 viral gene product varied. Some of the baits had less than five high-confidence host binding partners, such as NSP2, NSP4, NSP8, NSP9, NSP11, NSP12, NSP13, NSP14, NSP16, ORF3b, ORF7a, and ORF10, indicating that the major functions of these viral proteins may be carried out independently, with other viral proteins, and/or with very limited host functional protein partners. However, viral proteins N, ORF7b, M, and NSP6 had 102, 63, 53, and 48 host protein partners, respectively (**Figure 1H**). These viral gene products may work closely with host proteins to facilitate the viral lifecycle.

The fold change calculated by SAINTexpress analysis and the averaged peptide-spectrum matches (PSMs) of the identified preys are plotted in **Figure 1I**. The orange and red dots in the upper right represent HCIPs. The most significant interaction pair, ORF9b-TOMM70, was identified in more than 1000 PSMs of the prey. This interaction was reported recently in a structural study of the complex^6^. We also identified several binding pairs with very high MS signals, including ORF3a-VSP39, ORF3a-VSP11, ORF3a-CLCC1, NSP1-PYCR1, NSP1-PYCR2, N-G3BP1, and N-G3BP2. These strong interacting partners may have cellular functions important to SARS-CoV-2. The function of these preys may be taken over by the virus to suppress host cell activities and facilitate viral infection, replication, and maturation.

### Analysis of the Interactions between SARS-CoV-2 and Human Proteins

We next built an interaction network using the 437 identified virus-host protein-protein interactions. We found that many pathways or function complexes were highly enriched for binding to a specific SARS-CoV-2 gene product, as shown in **Figure 2**. Function enrichment may help us elucidate the biological functions and underlying mechanisms of these SARS-CoV-2 viral genes and design drugs targeting these virus-host interactions and/or the host complexes/pathways that are most relevant to this virus.

**Figure 2.**
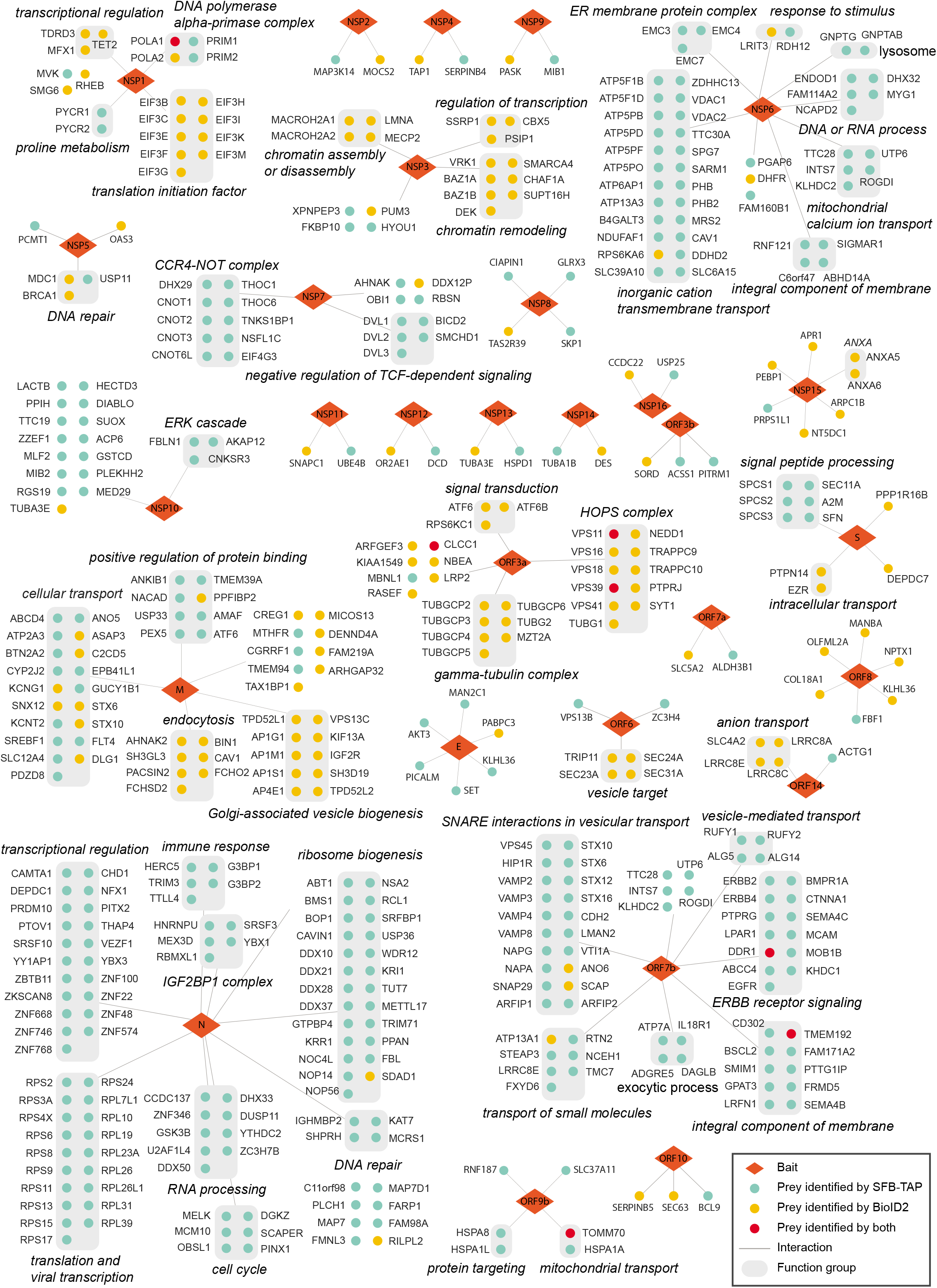
Interactomes between SARS-CoV-2 and human proteins. A protein-protein interaction network between 29 viral proteins of SARS-CoV-2 and the host proteins is shown for 437 pairs of host-virus interactions. The red diamonds are virus proteins (NSP, non-structure protein), green circles are the high-confidence interacting proteins (HCIPs) identified from SFB-tandem affinity purification experiments, yellow circles are the HCIPs identified from BioID2 labeling assays, and grey rounded rectangles are groups of proteins belonging to the same protein complex or from the same pathway. The italic text are the functional characterizations analyzed with Metascape and reference mining.

### SARS-CoV-2 Interacts with Host Proteins, which may Facilitate Viral Infection, Trafficking, and Budding

Most enveloped viruses do not encode their own membrane fission machinery, which is required for viral entry, transport, and budding^7^. Therefore, many host membrane proteins are recruited to work with the virus gene products to complete these functions. Our analysis of the SARS-CoV-2 virus-host protein-protein interaction network indicated that several viral gene products, such as M protein, NSP6, ORF3a, ORF6, and ORF7b, are involved in these processes.

M protein shares some host binding partners with another SARS-CoV-2 protein, ORF7b (**Figure 2**). Both proteins interact with several components of the SNARE complex, such as STX6 and STX10. The use of ORF7b as the bait led to the identification of more SNARE complex components, including STX12, STX16, VAMP2, VAMP3, VAMP4, VAMP8, NAPA, NAPG, and SNAP29. The SNARE complex is involved in eukaryotic membrane fusion and vesicular transport. This complex is known to be targeted by viruses during early infection, late viral production, and viral exit^8,9^. Many components of the HOPS complex, including VPS11, VPS16, VPS18, VPS39, and VPS41, were identified in interactions with the viral protein ORF3a (**Figure 2**). This complex plays a role in vesicle-mediated protein trafficking to lysosomal compartments and mediates tethering and docking events during SNARE-mediated membrane fusion through interaction with the SNARE complex^10^. The interactions of different viral proteins with distinct cellular complexes suggest that these associations are coordinated by the virus to maximize viral protein trafficking between different membrane-associated compartments.

CAV1 is well studied, with known functions in endocytosis and subsequent cytoplasmic transportation. Many steps during viral infection, such as the uptake of the virus, viral protein expression, and virion release, require CAV1 (Hoffmann and Pohlmann, 2018; Xing et al., 2020). The M protein and NSP6 of SARS-CoV-2 interact with CAV1 (**Figure 2**), suggesting that SARS-CoV-2 may involve CAV1 in both viral entry and a later step of the viral lifecycle.

Additionally, ORF3a interacts with TRAPPC9 and TRAPPC10 (**Figure 2**), subunits of the trafficking protein particle complex, which is known to have a function in vesicular transport from the endoplasmic reticulum to the Golgi apparatus. ORF6 binds to several components of the COPII complex (**Figure 2**), including SEC23A, SEC24A, and SEC31A, and may be involved in the formation of transport vesicles from the endoplasmic reticulum^11^. These host transportation systems are likely recruited by SARS-CoV-2 to facilitate its own gene expression, production, and transportation following viral infection.

We also identified a strong protein-protein interaction between ORF3a and CLCC1 (**Figure 2**). CLCC1 is a chloride ion channel protein with limited studies, and its functions are not well known. We obtained it as one of the top SARS-CoV-2 and host interaction proteins. In **Figure 3A**, we confirmed the interactions of ORF3a with CLCC1 and VPS11. These host proteins bind specifically to ORF3a. Furthermore, we examined the localization of CLCC1 with or without ORF3a overexpression (**Figure 3B**). CLCC1 is characterized as an endoplasmic reticulum protein, which was confirmed by our immunofluorescence results in cells without ORF3a overexpression. ORF3a is reported to be an endosome and lysosome-associated protein^12^, which was also validated by our immunofluorescence results. When ORF3a was overexpressed, the localization of CLCC1 changed to strong co-localization with ORF3a in the endosome and lysosome. Early reports suggest that chloride channel function may be required for the viral lifecycle^13,14^. Our results indicate that SARS-CoV-2 may take over the function of CLCC1 protein to promote its own lifecycle. Although the detailed mechanisms remain to be elucidated with future functional analysis, the strong interaction between ORF3a and CLCC1 indicates that disrupting this interaction and/or CLCC1 function may affect the SARS-CoV-2 lifecycle.

**Figure 3.**
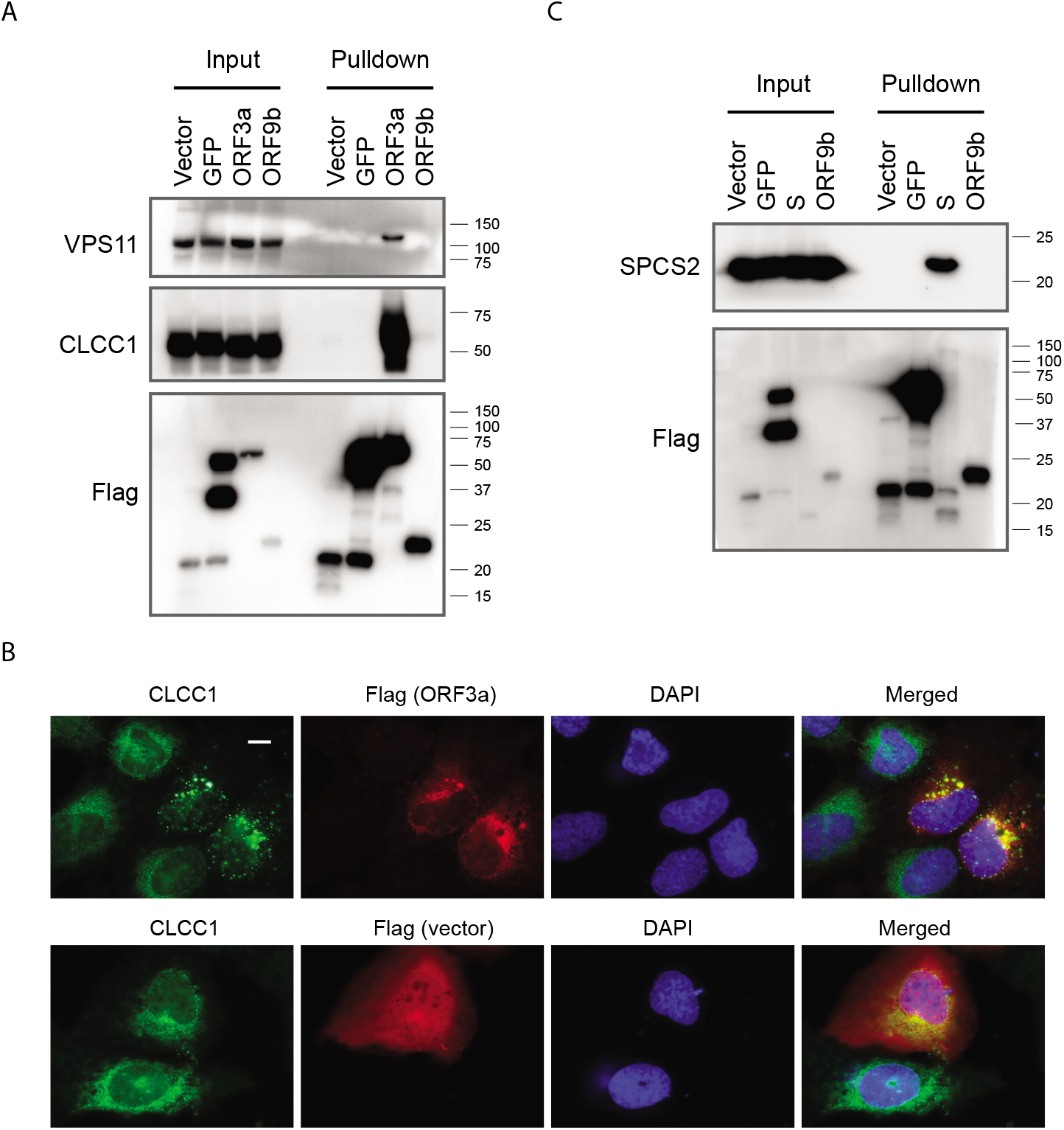
Validation of selected host-virus protein-protein interactions. (A) Pulldown and Western blot analysis validated the interaction between the viral protein ORF3a and its interactors, VPS11 and CLCC1. HEK293T cells with SFB-tagged ORF3a expression were collected and lysed. Cell lysates were subjected to pulldown assay using S-protein beads. Western blot analysis was conducted with the indicated antibodies. Cells transfected with vector or construct encoding GFP or ORF9b were included as controls in these experiments. (B) Immunostaining analysis of protein localization. U2OS cells were transfected with construct encoding ORF3a. Cells were fixed and stained with the indicated antibodies. The green signal is CLCC1, the red signal is flag (for SFB-ORF3a), and the blue signal indicates DAPI/nuclei. Scale bar: 10 μm. (C) Pulldown and Western blot validation of the interaction between SARS-CoV-2 S protein and the human signal peptidase complex. Cells transfected with vector or construct encoding GFP or ORF9b were included as controls in these experiments.

In summary, we identified several SARS-CoV-2 gene products, M protein, NSP6, ORF3a, ORF6, and ORF7b, that interacted with host cell membrane proteins and complexes. The transmembrane domain prediction by TMHMM 2.0 also suggested that these viral gene products have at least one transmembrane domain in their protein sequences, except for ORF6, which is a short protein with only 61 amino acids. These viral proteins may have key functions and recruit host membrane proteins and complexes to facilitate viral infection, trafficking, and/or budding.

### SARS-CoV-2 Manipulates the Host Cell Replication, Transcription, and Translation Machinery

After a virus infects a host cell, it needs to suppress the host cell replication, transcription, and translation processes and recruit those factors for the lifecycle of the virus. According to our virus-host interactome analysis, NSP1, NSP3, NSP5, and N protein of SARS-CoV-2 are the major viral proteins involved in the regulation of host replication, transcription, and translation.

NSP1 is a well-studied coronavirus non-structural gene product that displays biological functions in suppressing host gene expression^15–17^. Accordingly, we identified nine HCIPs of eukaryotic initiation factor 3 (EIF3) complex subunits that bind to NSP1 (**Figure 2**). EIF3 is the largest translation initiation factor complex, composed of 13 subunits. It is involved in nearly all steps of translation initiation^18^. The EIF3 complex has been shown to be a virus binding complex that results in the suppression of host gene expression^19^, which is consistent with our interactome results.

In addition to the HCIPs functioning in translation regulation, we also identified the whole DNA polymerase alpha complex, i.e., POLA1, POLA2, PRIM1, and PRIM2, in the NSP1 interactome (**Figure 2**), which is similar to a recent report^3^. Moreover, our interactome analysis revealed that NSP5 associates with several well-studied DNA damage proteins, MDC1, BRCA1, and USP11 (**Figure 2**). NSP1 and NSP5 may interact with these host replication and/or DNA damage-related proteins to inhibit the initiation of host genome replication and DNA damage response and/or facilitate viral genome replication.

NSP3 is the largest gene product of coronaviruses, with multiple functional domains that may be involved in the regulation of replication and/or transcription. The interactome of NSP3 has not been well studied owing to the large protein size of NSP3. In our interactome analysis, we expressed the full length of NSP3 in both SFB-TAP and BioID2 labeling experiments and successfully identified >100 PSMs for the bait NSP3. After HCIP analysis, we found that NSP3 interacts with several host proteins involved in transcription regulation as well as in chromatin assembly and remodeling (**Figure 2**). BAZ1A, BAZ1B, and SMARCA4 function in the ISWI complex to control DNA accessibility and chromatin remodeling^20^ and may have a role in transcription regulation during SARS-CoV-2 infection. MACROH2A1, MACROH2A2, SSRP1, and SUPT16H are nucleosome factors that may also control gene transcription by wrapping and compacting DNA into chromatin. With these and other preys that may also play roles in transcription regulation, we believe that the major function of NSP3 is transcription regulation, which may suppress host gene expression and/or facilitate viral gene expression by binding to and interfering with multiple host proteins involved in transcription or chromatin control.

Another viral transcription and translation regulator we identified in the interactome analysis is the N protein. N protein is a nucleocapsid protein that binds directly to the virus genome. We identified many ribosome proteins with strong interaction with N protein (**Figure 2**). This is similar to reports in other virus species showing that viral N proteins bind to host ribosome subunits^21,22^. Ribosome proteins are required for viral gene translation and viral genome replication^23^. SARS-CoV-2 N protein is the first released virus protein after the infection. It may bind to ribosome proteins quickly to facilitate the translation of other virus genes. At the next step, a group of ribosome proteins may be recruited by N protein to facilitate virus genome replication^24^. Our data also indicate that the N protein binds to the IGF2BP1 complex, which has been shown previously to enhance viral translation initiation and stabilize viral RNA^25^. In addition to the above-mentioned HCIPs, we identified many other RNA processing, RNA metabolism, and transcriptional regulatory proteins as N protein interacting partners (**Figure 2**). These data strongly suggest that N protein is one of the major regulators of SARS-CoV-2 involved in viral transcription, translation, and genome replication.

We also showed that the signal peptidase complex, including SPCS1, SPCS2, SPCS3, and SEC11A, significantly binds to the S protein (**Figure 2**). The signal peptidase complex is required for post-translational processing of proteins involved in virion assembly and secretion^26–28^. This complex may also promote the interaction between virus structure proteins, which benefits virus assembly^28^. Indeed, we confirmed the interaction of S protein with the signal peptidase component SPCS2 (**Figure 3C**), suggesting that the S protein of SARS-CoV-2 may function similarly to S proteins in other viruses.

### SARS-CoV-2 Genome Replication Can Be Mediated through Interactions with Several Host Proteins

Three components of endoplasmic reticulum membrane protein complex, EMC3, EMC4, and EMC7, were identified as NSP6 interacting proteins (**Figure 2**). Large-scale genetic screening revealed that EMC components may be the essential genes involved in viral replication and downstream cell death pathways^29^. Another genomic screen showed that EMC components are needed during the early stages of viral infection^30^. The interaction between NSP6 and multiple EMC components suggests that SARS-CoV-2 interacts with the EMC through NSP6, which may help viral infection and replication.

Moreover, the NSP7-NSP8 complex forms the primase that functions as RNA replicase. With overexpression of NSP7 alone in HEK293T cells, we captured a group of proteins belonging to the CCR4-NOT complex, including CNOT1, CNOT2, CNOT3, and CNOT6L (**Figure 2**). The CCR4-NOT complex functions in mRNA decay by shorting poly(A) and is associated with translating ribosomes and RNA processing bodies. A genome-wide genetic screen to study hepatitis C virus pathogenesis revealed that these four genes are required for viral infection^31^. The CCR4-NOT complex may facilitate viral replication by triggering the turnover of nucleoprotein-bound cellular mRNA and releasing the nucleoprotein to bind viral genomic RNA for replication^32^. With a similar mechanism, the interaction between NSP7 and the CCR4-NOT complex may facilitate replication of the SARS-CoV-2 genome.

Another replication function–related complex we identified as a HCIP of NSP7 was the THO/TREX complex. We identified THOC1 and THOC6 as NSP7-interacting proteins (**Figure 2**). The THO/TREX complex mediates transcription elongation and nuclear export. A study of a Kaposi sarcoma-associated herpesvirus suggests that this complex is essential for stabilization and export of viral intron-less mRNAs and production of infectious viral particles^33,34^. Interaction between the viral gene and the THO/TREX complex occurs in an ATP-dependent manner, which suggests potential drugs to target this interaction^35^. Similar drugs may also help in the treatment of COVID-19.

Taken together, our results showed that NSP6 and NSP7 of SARS-CoV-2 interact with several protein complexes, and these virus-host protein-protein interactions may inhibit host functions and facilitate viral genome replication and virus production.

### SARS-CoV-2 Controls the Host Cell Metabolism and Signaling Pathways

Mitochondria are the major targets manipulated by the virus after infection. In taking control of the mitochondria, the virus disrupts host cell functions and forces cells to produce energy and other products needed for the viral lifecycle^36,37^. NSP6 was predicted with eight transmembrane helices when analyzed with TMHMM 2.0. The SARS-CoV-2 gene NSP6 may play a major role in hijacking the mitochondria. Interactome analysis of NSP6 revealed that it interacted with a few subunits of ATP synthase (**Figure 2**), e.g., ATP5F1B, ATP5F1D, ATP5PB, ATP5PF, ATP5PO, ATP6AP1, and ATP13A3, indicating that the virus may interfere directly with the machinery involved in cellular energy production. The interaction of NSP6 with VDAC1 and VDAC2 may enhance viral infection and replication, as reported in a study of an infectious bursal disease virus that induces cell death^38,39^. NSP6 also binds to PHB and PHB2, which are components of a large ring complex, the prohibitin complex, which stabilizes mitochondrial respiratory enzymes and maintains mitochondrial integrity^40^. These interactions of NSP6 with mitochondrial proteins suggest that NSP6 may manipulate the host energy production system to serve the lifecycle of SARS-CoV-2.

NSP6 may also participate in signal transduction; we identified SIGMAR1 as an important binding protein of NSP6. SIGMAR1 is an endoplasmic reticulum membrane protein that plays a role in lipid transport and calcium signaling. The interaction between NSP6 and SIGMAR1 has been studied recently^3^, and an antiviral strategy against this prey was found to be potentially effective. Another SARS-CoV-2 gene involved in signaling pathways was NSP7. We identified DVL1, DVL2, and DVL3 as NSP7 interacting proteins (**Figure 2**). These proteins are central components of the Wnt/beta-catenin signaling pathway. Previous studies revealed that activation of the Wnt/beta-catenin signaling pathway could stimulate viral replication^41,42^, and inhibition of this pathway could reduce the viral yield during productive viral infection^43,44^. Moreover, we showed that NSP10 interacts with several factors involved in the ERK1/ERK2 signaling pathway (**Figure 2**), implying that SARS-CoV-2 may hijack the MAPK-ERK pathway to control the cell fate for the viral lifecycle, as reported for other viruses^45^.

### SARS-CoV-2 Fights Host Cell Immune and Antiviral Systems

Host cells have developed efficient antiviral systems to counter viral infections. Similarly, viruses find ways to inhibit host antiviral systems to ensure survival. The interferon (IFN)-mediated antiviral pathway is one of the major innate immune responses against viral pathogens. We expressed the 29 SARS-CoV-2 coding genes in HEK293T cells and then analyzed the activity of IFN-beta luciferase reporter response to infection with Sendai virus, or the activity of ISRE-luciferase reporter response when viral proteins were co-expressed with IRF3 (**Figure S2**). We found that two viral proteins, NSP1 and N protein, most significantly suppressed the activation of IFN signaling based on these two assays.

NSP1 acts as the strongest gene product in SARS-CoV-2 that helps evade host cell antiviral defense. It suppresses global gene expression in the host as we mentioned above. Our host-virus interactome results showed that NSP1 interacts with nine components of the EIF3 complex to suppress the translation machinery. Translation of host antiviral genes is also inhibited by this global translation suppression function. Therefore, NSP1 acts as the strongest viral protein that inhibits IFN-beta signaling when overexpressed.

N protein is the first released and most abundant SARS-CoV-2 viral protein upon infection. We identified several immune response proteins that are recruited by N protein (**Figure 2**), and these proteins may facilitate the viral infection and later the viral lifecycle. Similar to a previous study^3^, we identified strong interactions between N protein and Ras-GTPase-activating protein-binding protein 1/2 (G3BP1/2; **Figure 2**). G3BP1 plays essential roles in innate immunity by activating the cGAS/STING pathway and acting as a platform for antiviral signaling^46^. In a previous study of an RNA virus, G3BP1 was found to enlist an antiviral response by positively regulating RIG-I^47^. The strong binding of G3BP1 and G3BP2 with N protein may inhibit cGAS activation when the SARS-CoV-2 genome is released into host cells. Another antiviral protein, HERC5, was also identified as an N protein interactor (**Figure 2**). HERC5 is the major E3 ligase for ISG15 conjugation, and ISGylation of IRF3 is required to sustain its activation and boost the antiviral response^48,49^. N protein also interacts with TTLL4 (**Figure 2**), which regulates the nucleotidyl-transferase activity of cGAS. Thus, the N protein likely represents the first wave of suppressors of host antiviral mechanisms to ensure the survival of SARS-CoV-2 upon infection.

In addition, NSP5, which is the 3C-like protease of SARS-CoV-2, cleaves the ORF1a/ORF1ab polyprotein into function fragments. Our interactome analysis revealed a potential interaction between NSP5 and an IFN-induced antiviral enzyme, OAS3 (**Figure 2**). OAS3 is an IFN-induced antiviral gene that plays a critical role in cellular innate antiviral response^50,51^. OAS3 is activated by dsRNA generated after viral infection, and the enzyme then leads to the activation of RNase L to digest the viral RNA and terminate its replication. We speculate that NSP5 may bind to and cleave OAS3 to inhibit its antiviral function.

We also found that both M protein and ORF7 interact with growth factor receptors (**Figure 2**). M protein interacts with FLT4 (VEGFR3), and M protein may be involved in a selfcontrol mechanism to prevent uncontrolled inflammation in macrophages during bacterial infection^52^. A group of ORF7-binding proteins are ERBB receptors and proteins involved in related signaling pathways, including EGFR (ERBB1), ERBB2, and ERBB4 (**Figure 2**). There are four ERBB receptors, and these are suggested to function as gatekeepers of infectious diseases^53^. EGFR was reported to help viruses enter into host cells^54–56^. Similarly, ERBB2 was also targeted by pathogens for cellular entry^57^. The membrane association of EGFR and ERBB2 was increased during viral infection to benefit the processing of hepatitis C virus in previous studies^58,59^. Moreover, ERBB receptors are potent factors promoting cellular growth and survival, which could prolong host cell survival^60,61^ and modulate host immune responses^62,63^ to benefit viral production. SARS-CoV-2 is suspected to bind to growth factor receptors and regulate the related signaling pathways to suppress host antiviral systems and promote host cell survival to increase viral infection and production.

M protein also interacts with several subunits of clathrin-associated adaptor protein complexes, AP1G1, AP1M1, AP1S1, and AP4E1 (**Figure 2**). These adaptor proteins are involved in protein sorting, vesicle formation, and cargo selection. Binding between adaptor protein complex 1 and BST2, an antiviral restriction factor that inhibits the release of enveloped viruses, was shown to retain BST2 in endosomes and stimulate its degradation^64^. The interaction between M protein and these clathrin-associated adapter protein complexes may suggest mechanisms that lead to mis-trafficking and degradation of host proteins to facilitate viral infection and the viral lifecycle.

Collectively, the association between several viral proteins and host proteins involved in antiviral defense and other cellular functions may indicate that SARS-CoV-2 employs multiple ways to evade host antiviral systems for its survival, reproduction, and release.

### Potential Drug Targets Revealed by the Host-Virus Interactome Study

The interactions between viral and host proteins are believed to play fundamental roles in the virus lifecycle. It is reasonable to speculate that drugs targeting host proteins may inhibit the functions of host protein complexes and/or disrupt their interactions with viral proteins, which may lead to failure of the virus lifecycle. Therefore, we analyzed potential drug targets and drugs using bioinformatics tools, Metascape^65^ or Ingenuity Pathway Analysis. Of the 29 viral proteins and 437 SARS-CoV-2 and host HCIPs, we found 18 viral proteins with 60 host interaction proteins that could be targeted by various drugs (**Figure 4 and Table S3**). Of these potentially drugs targets, some well-studied drugs are available and/or being developed. SIGMAR1, which binds to NSP6, was tested as an important drug target candidate for the treatment of COVID-19^3^. Many of our candidate proteins are well studied as drug targets, such as DHFR and VDAC1/VDAC2, which interact with NSP6; GSK3B, which interacts with N protein; FLT4, which interacts with M protein; EGFR, ERBB2, and ERBB4, which interact with ORF7b; A2M, which interacts with S protein; and ANXA5, which interacts with NSP15. Multiple drugs targeting these host proteins have been developed, which should facilitate the testing and developing treatments for COVID-19 patients. Two potential drugs proposed in our study were tested with real-world COVID-19 treatment, haloperidol (SIGMAR1 antagonist) and tamoxifer (ERBB2/ERBB4 inhibitor) (**Figure 4**). And they both showed effectiveness against COVID-19^6^.

**Figure 4.**
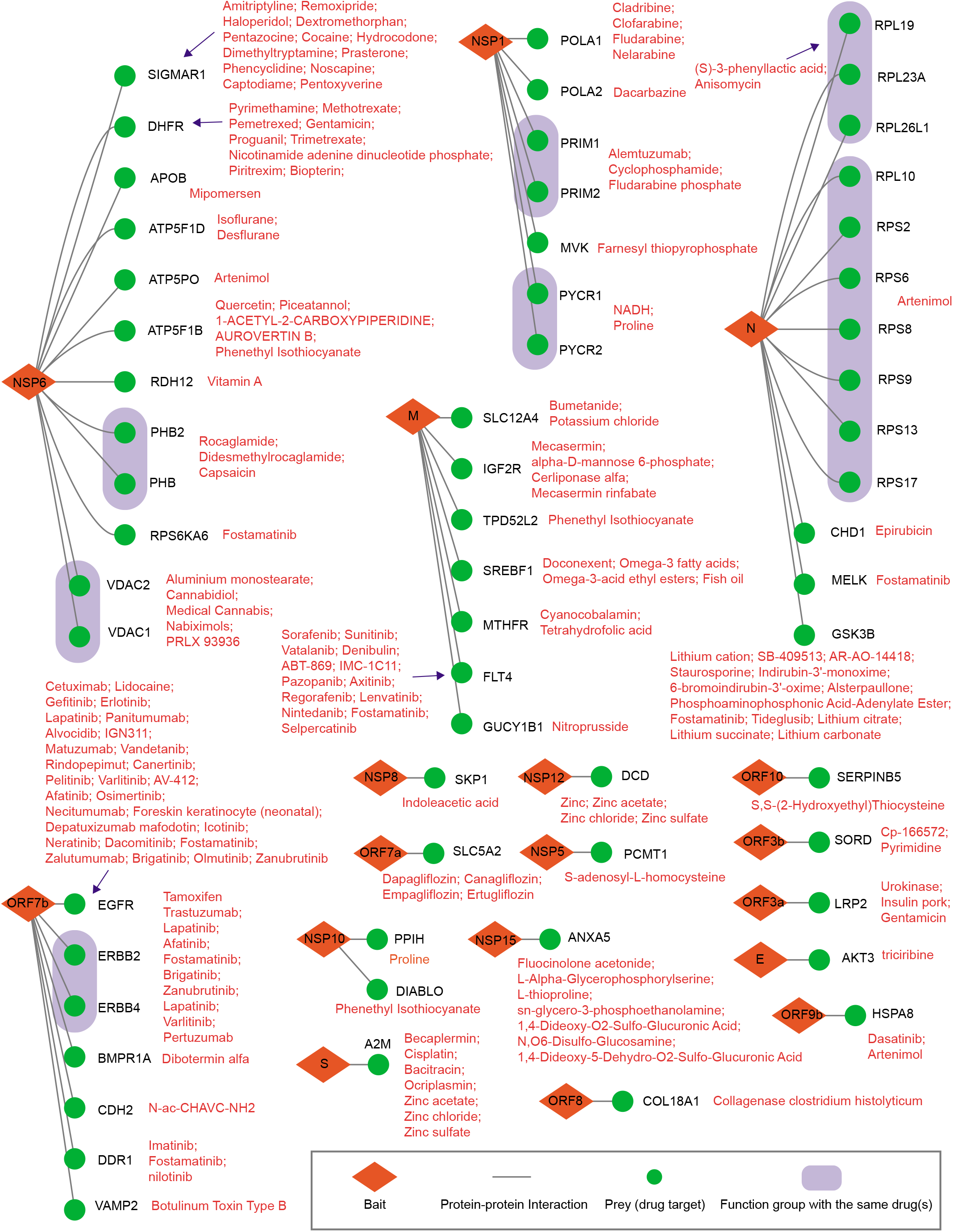
Potential drugs targeting SARS-CoV-2 and host protein interaction. The drug-human target network was analyzed by Metascape and Ingenuity Pathway Analysis/GO analysis. The red diamonds are virus proteins, green circles are the high-confidence interactive proteins that were identified as drug targets, and red text lists potential drugs. Proteins in the blue rounded rectangles belong to the complex or functional pathway targeted by the same drugs.

### Comparison of Coronavirus-Host Interaction Networks among Human Coronaviruses

Seven human coronaviruses have been reported since the first one, HCoV-OC43, was identified in the mid-1960s. Four of these coronaviruses cause mild to moderate symptoms, including HCoV-OC43, HCoV-HKU1, HCoV-NL63, and HCoV-229E. The other three coronaviruses, SARS-CoV-2, SARS-CoV-1, and MERS-CoV, can cause more serious, even fatal diseases. To understand the overlap and divergence of virus-host interactions among these human coronaviruses, we selected two viral gene products, NSP1 and N protein, for comparison. These two viral proteins also significantly suppress the host antiviral IFN pathway as shown in **Figure S2**. These comparisons were meant to identify potential targets that could be used for pan-viral treatment against currently known coronaviruses and any new coronaviruses identified in the future. The comparisons may also reveal different mechanisms adopted by each coronavirus for its viral lifecycle and associated pathologic symptoms.

### Comparison of NSP1 Interactomes among All Seven Human Coronaviruses

NSP1 is arguably the most important pathogenic determinant of human coronaviruses, because it induces a near-complete suppression of host gene expression^66^. NSP1 uses a two-pronged strategy for this host gene suppression, i.e., stalling canonical mRNA translation and triggering the degradation of host mRNAs^67^. NSP1 also suppresses host antiviral gene expression, which inhibits the host antiviral defense systems and therefore facilitates viral replication and immune evasion^68^. NSP1 proteins from two alpha coronaviruses, CoV-229E and CoV-NL63, are shorter than those from the other five beta coronaviruses (**Figure 5A**). Most coronavirus NSP1 proteins share low sequence identity, i.e., 10% to 20%, except for SARS-CoV-2 and SARS-CoV-1, which have a shared sequence identity of 84.4% (**Figure S3A**). However, all of these NSP1 proteins have similar functions in suppressing host gene expression and inhibiting the host antiviral system, such as IFN signaling (**Figures S3B and S3C**).

**Figure 5.**
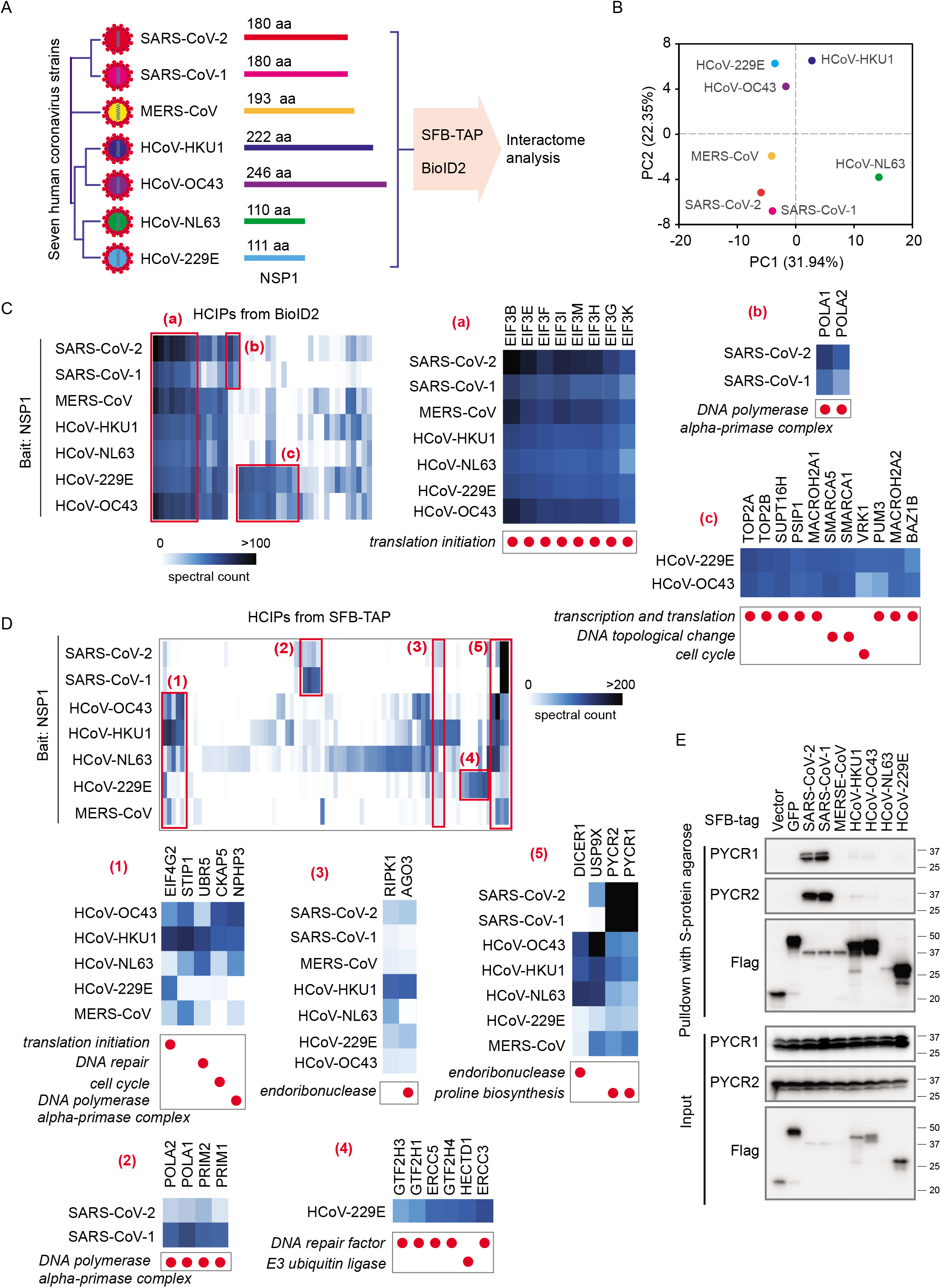
Comparison of interactomes of seven NSP1 proteins from seven human coronaviruses. (A) Comparisons of NSP1 proteins from seven human coronaviruses. (B) Principal component analysis of the seven HCIP lists with bait NSP1 from different human coronaviruses. (C, D) Heat map of the NSP1 HCIPs obtained using (C) BioID2 and (D) SFB-TAP. NSP1 HCIPs were compared among seven human coronaviruses using the preys’ spectral counts. Three areas of the heat map in (C) are enlarged and labeled as (a), (b), and (c). Five areas of the heat map in (D) are enlarged and labeled as (1), (2), (3), (4), and (5). Functional characterization of each protein is shown with the red circles below the enlarged images. (E) Pulldown and Western blot validation of the interaction between human coronavirus NSP1 proteins and the human proteins PYCR1/PYCR2. Cells transfected with vector or construct encoding GFP were included as controls in these experiments.

We expressed all seven human coronavirus NSP1 proteins in HEK293T cells and performed both SFB-TAP (**Figure S3D**) and BioID2 labeling (**Figures S3E and S3F**) experiments to uncover the protein-protein interaction networks between NSP1 and host proteins, following a similar strategy as described in the SARS-CoV-2 interactome analysis presented in **Figure 1**. To compare all HCIPs among different human coronavirus strains, we first identified proteins for each coronavirus NSP1 with a SAINTexpress Bayesian false discovery rate ≤0.01 in at least one of the seven coronavirus NSP1 interactions. Using these HCIPs, we identified all PSMs captured for that prey in the raw identification data for each coronavirus NSP1 (**Table S4 and S5**). Principal component analysis grouped SARS-CoV-2 and SARS-CoV-1 together (**Figure 5B**), indicating that these two viruses with high NSP1 sequence identity share similar interactomes and may adopt similar mechanisms to suppress host gene expression. The protein-protein interaction network of NSP1 in the seven human coronaviruses was built in **Figure S3G**.

Although many of these NSP1 proteins have limited sequence identity, we uncovered subsets of shared HCIPs across all seven human coronavirus NSP1 interactomes. In BioID2-MS analysis, we identified the EIF3 complex, including eight of the thirteen subunits in our NSP1 HCIP list (**Figure 5C**). The EIF3 complex stimulates nearly all steps of translation initiation and is essential for most forms of cap-dependent and cap-independent translation initiation^18^. In our TAP-MS analysis, a translation initiation factor, EIF4G2, was identified in all seven coronavirus NSP1 interactomes (**Figure 5D**). When the cell is infected by coronavirus, the viral NSP1 protein may recognize translation initiation factors and suppress almost all translation activities of the host cell, which may be similar to the situations reported for human immunodeficiency virus and other important human pathogens^19^. Additionally, several other transcription and translation regulators, such as BCLAF1, TDRD3, SSRP1, SMG6, and DIS3L2, were identified across all seven coronavirus NSP1 interactomes. The analysis of common NSP1-binding partners across all seven coronaviruses may explain why NSP1 proteins with only 10% to 20% sequence identity have similar functions in suppressing host gene expression. In addition, these common HCIPs may represent potential drug targets against all human coronaviruses, including any identified in the future.

In contrast to the common HCIPs for all coronaviruses, some HCIPs were unique or showed significantly higher binding signals to NSP1 from one or several coronaviruses. For example, the subunits of DNA polymerase alpha-primase complex, POLA1, POLA2, PRIM1, and PRIM2, were identified with NSP1 of SARS-CoV-1 and SARS-CoV-2, which was reported recently^6^. However, there was no or weak signal of this complex in other coronavirus NSP1 interactomes (**Figures 5C and 5D**). We identified more than 500 PSMs for both PYCR1 and PYCR2 in SARS-CoV-2 and SARS-CoV-1 purifications (**Figure 5D**), indicating that these two viruses have some unique mechanisms. We further confirmed the SARS-CoV-2 and SARS-CoV-1 NSP1-PYCR1/PYCR2 protein-protein interaction by immunoblotting (**Figure 5E**). PYCR1/PYCR2 proteins catalyze the last step in proline biosynthesis. This process is critical for tumorigenesis^69,70^ and may be hijacked by human herpesvirus for tumorigenesis^71^. The functional significance of NSP1-PYCR1/PYCR2 interaction in viral lifecycle warrants further investigation.

Additionally, one group of proteins was identified only in HCoV-229E and HCoV-OC43 NSP1 purifications (**Figure 5C**); these proteins include the DNA topoisomerase subunits TOP2A and TOP2B, global transcription activators SMARCA1 and SMARCA5, and other transcription or translation regulators (SUPT16H, PSIP1, MACROH2A1, MACROH2A2, PUM3, and BAZ1B). An RNA-mediated gene silencing component, AGO3, was identified in several coronaviruses, but with much higher signal in HCoV-HKU1 (**Figure 5D**). Another protein, RIPK1, which is a key regulator of tumor necrosis factor-mediated apoptosis, necroptosis, and inflammatory pathways, was also identified as binding with all seven human coronavirus NSP1 proteins, but the strongest interaction was in HCoV-HKU1. Both AGO3 and RIPK1 play a role in NF-kappaB signaling, which may regulate NF-kappaB activation, constituting a major antiviral immune pathway^72^. A general transcription and DNA repair factor, IIH core complex, was identified as uniquely binding to HCoV-229E; we identified several of its subunits, including GTF2H1, GTF2H2, GTF2H4, ERCC3, and ERCC5 (**Figure 5D**). DICER1, which plays a central role in post-transcriptional gene silencing and exogenous RNA degradation, was identified with significantly higher PSMs in HCoV-OC43, HCoV-HKU1, and HCoV-NL63 than in the other coronaviruses (**Figure 5D**). Together, these findings suggest that different and unique mechanisms may be utilized by each coronavirus after infection to suppress host gene expression. These differences in NSP1 interactomes may also explain why different coronaviruses have different symptoms in humans.

### Comparison of N Protein Interactomes among Human Coronaviruses

SARS-CoV-2 N protein is the first released and most abundant protein during viral infection. N protein has several key functions that help viral transcription, translation, genome replication, and packaging before budding. N protein is also one of the key molecules that the virus uses to fight against host antiviral systems. We investigated the interactomes of N protein in different coronaviruses using the same strategy as described above for the study of NSP1 (**Figure 6A**). Most of the N proteins in different human coronaviruses had sequence identity percentages between 20% and 30%, but the sequence identity was 89.1% between SARS-CoV-2 and SARS-CoV-1 (**Figure S4A**).

**Figure 6.**
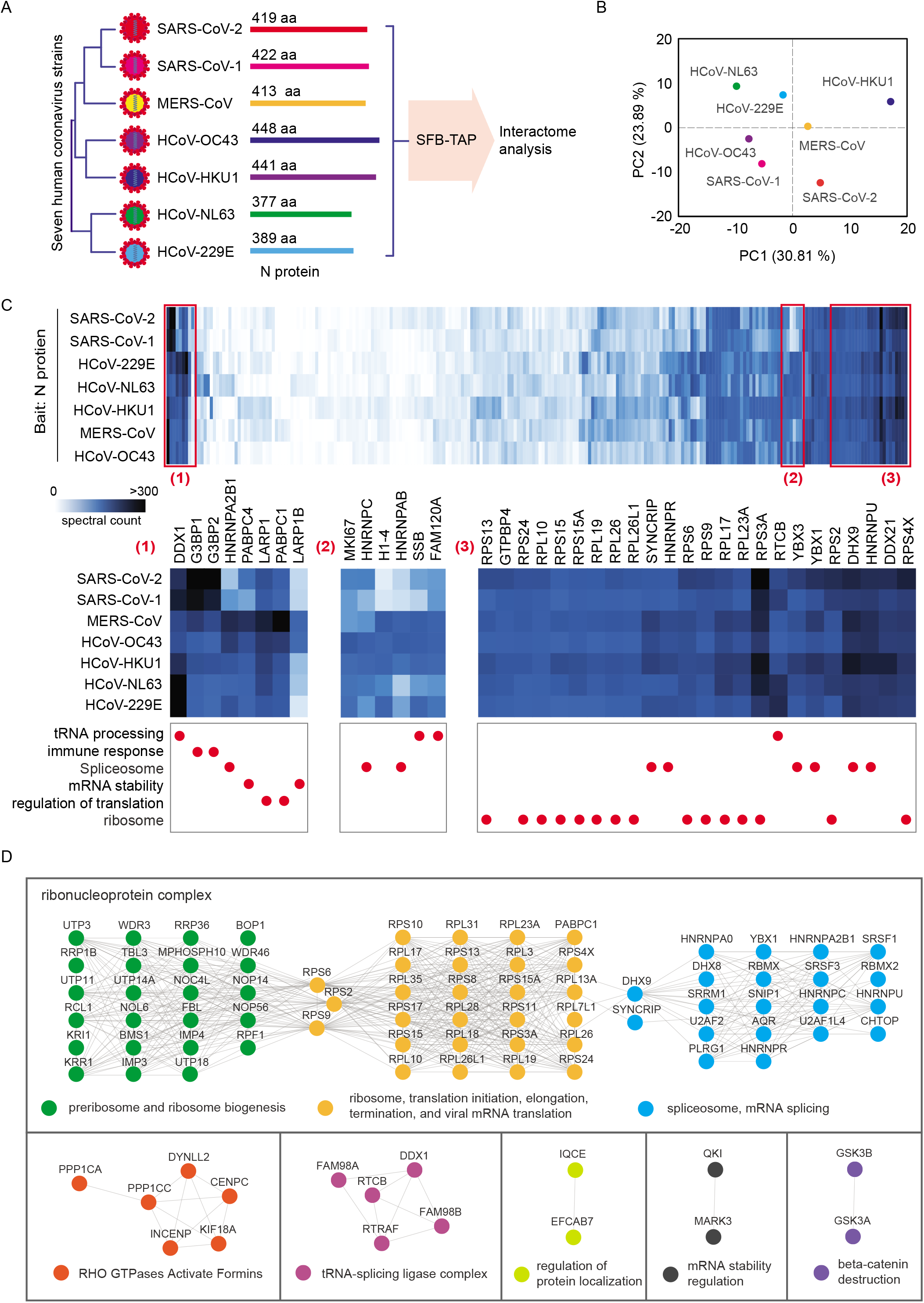
Comparison of interactomes of N proteins from different human coronaviruses. (A) Comparison of N proteins from seven human coronaviruses. (B) Principal component analysis of the seven high-confidence interacting protein (HCIP) lists with bait N protein from different human coronaviruses. (C) Heat map of the N protein HCIPs identified by SFB-tandem affinity purification. N protein HCIPs were compared among seven human coronaviruses using the preys’ spectral counts. Three areas of the heat map are enlarged and labeled as (1), (2), and (3). Functional characterization for each protein is shown with the red circles below the enlarged images. (D) The human proteins which binding to N proteins of human coronaviruses were integrated and analyzed by STRING to illustrate the protein-protein interaction among the HCIPs. The minimum required interaction score setting in STRING is 0.9. Different function group or complex were colored differently. The cycles are proteins, and the lines are the interactions.

In the interactome analysis of all SARS-CoV-2 coding genes, we identified the most HCIPs with N protein as bait, as shown in **Figure 2**. Among the 102 HCIPs identified, we found that 100 of them were identified by SFB-TAP. Therefore, we focused on SFB-TAP for all seven human coronaviruses. We expressed all seven human coronavirus N proteins in HEK293T cells and performed SFB-TAP (**Figure S4B**), and then we built the protein-protein interaction network of N proteins in the seven human coronaviruses (**Figure S4C**). Similar to the analysis of NSP1 interactomes, we performed a principal component analysis (**Figure 6B**) and generated a heat map of the N protein interactomes in different coronaviruses (**Figure 6C and Table S6**). Most of the strong binding HCIPs were shared among all coronaviruses, and a large portion of the shared HCIPs play a role in RNA processing, ribosome biogenesis, and translation regulation (**Figure 6D**).

We tested IFN signaling suppression by N proteins of all seven human coronaviruses. Our results indicated only SARS-CoV-2 and SARS-CoV-1 N proteins suppressed the activation of IFN-signaling significantly (**Figures 7A and 7B**). We analyzed N proteins interacting proteins, and found two differential interacting proteins in the N protein interactomes among human coronaviruses is G3BP1 and G3BP2. As shown in **Figures 6C and 7C**, we recovered around 400 PSMs for these two preys in the purification of SARS-CoV-2 N protein, which was several times higher than those in MERS-CoV and other four common human coronaviruses. We confirmed these bindings with immunoblotting (**Figure 7D**). G3BP1 and G3BP2 bind more strongly with the N proteins from SARS-CoV-2 and SARS-CoV-1 than with the N proteins from other human coronaviruses, and the binding to MERS N protein was the weakest. G3BP1 and G3BP2 are important for innate immunity, as described above. We suspect that the binding affinity between N proteins and G3BP1/G3BP2 may in part contribute to the inhibition of the host antiviral system and lead to differences in the contagiousness of these viral pathogens.

**Figure 7.**
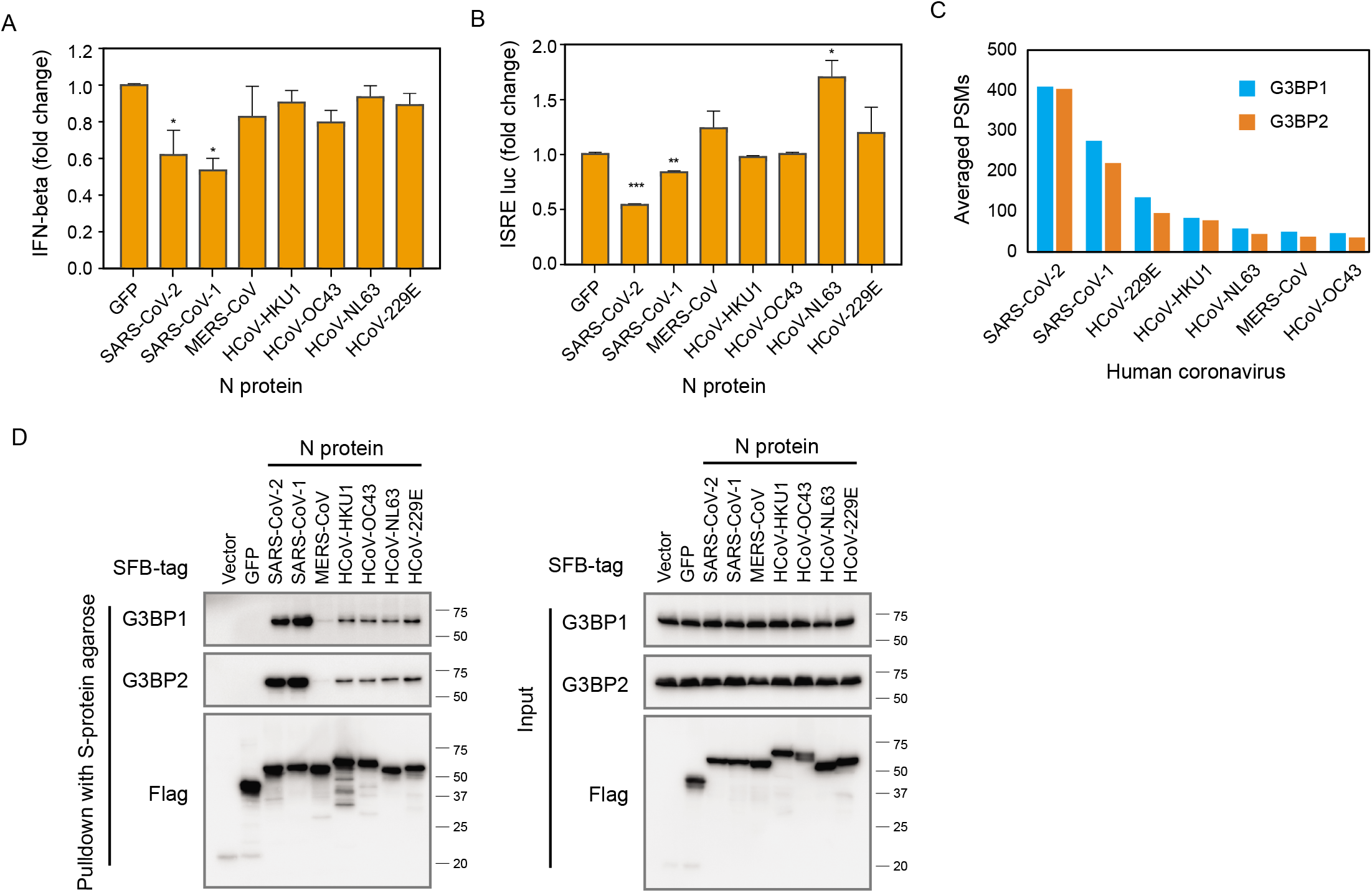
SARS-CoV-2 N protein interacts with G3BP1/G3BP2, which may in part contribute to fighting the host antiviral system. (A, B) Interferon (IFN) signaling assay for N proteins of seven human coronaviruses. (A) Analysis of IFN-beta-luciferase reporter 12 hours after Sendai virus infection. (B) Analysis of ISRE luciferase reporter when the viral proteins were co-expressed with IRF3. Graphs show mean ± SD, n = 3. ***p < 0.001, **p < 0.01 and *p < 0.05 (Student’s *t*-test). (C) Spectra counts of G3BP1/G3BP2 identification in purifications with N proteins from different human coronaviruses as baits. PSM, peptide-spectrum match. (D) Pulldown and Western blot validation of the interaction between N protein and G3BP1/G3BP2. Cells transfected with vector or construct encoding GFP or N protein from different human coronaviruses were compared in these experiments.

## DISSUSION

In the current study, we generated a comprehensive host-virus protein-protein interaction network of SARS-CoV-2 by identifying HCIPs using two different interactome approaches, i.e., TAP-MS and BioID2-MS. We identified a total of 437 HCIPs that bind to one or several of the SARS-CoV-2 genes. These findings not only validate several known protein-protein interactions but also identify many new protein-protein interactions with potential functions in the SARS-CoV-2 lifecycle. This dataset reveals new mechanistic insights into the diverse functions of SARS-CoV-2 genes, which need to be further investigated. Moreover, this dataset suggests potential drug targets for the treatment of COVID-19.

We identified 314 HCIPs from TAP-MS experiments and 130 HCIPs using the BioID2-MS strategy. Surprisingly, we found that only seven proteins overlapped between these two methods, as shown in **Figure S5A**. When we examined the details of the HCIP lists, we found that the TAP and BioID2 methods have their own biases. The TAP method is effective in identifying HCIPs for soluble proteins, whereas BioID2 facilitates the detection of binding partners of baits with transmembrane domains, which is similar to previous studies using this method^4^. In our SARS-CoV-2 interactome results, we identified many HCIPs for three baits when using the BioID2 method. All three proteins are predicted to have multiple transmembrane domains according to our analysis using TMHMM, i.e., NSP3 (four domains), M (three domains), and ORF3a (three domains). However, for the baits with only one predicted or known transmembrane domain, such as S, E, and ORF7b protein, our TAP method worked efficiently.

Another possible explanation for the difference between the TAP and BioID2 methods may be influence from the data obtained and the subsequent data analysis. Because the TAP method employs TAP, whereas BioID2 uses one-step purification with streptavidin beads, we uncovered much shorter raw protein/peptide lists with the TAP-MS method, which allowed the MS instrument to capture much more PSMs for each HCIP. For example, the ORF9a–TOMM70 interaction was uncovered as an HCIP by both methods. However, we detected an average of 1518 PSMs of TOMM70 from three biological replicates in our TAP-MS experiments, but only an average of 31 PSMs in the BioID2-MS study. Our current data analysis of HCIPs may not be adequate to deal with this technical issue, because it is challenging to distinguish true binding partners from contaminants in a long raw protein/peptide list with few prey PSMs for each protein. The third explanation for differences between the two methods is the restriction of BioID2 or any other proximity-labeling method, which limits the coverage of binding partners located beyond the small labeling reaction range. The overlap between the interactomes revealed by different strategies is quite low^6,73^ (**Figure S4B**). This is still a common issue in protein-protein interaction studies, and it may due to the purification strategy, experimental design, and/or data filtration/analysis.

We built a host-virus protein-protein interaction network of SARS-CoV-2 based on the 437 HCIPs identified in the current study. It may suggest how these viral proteins participate in the virus lifecycle. M protein, NSP6, ORF3a, ORF6, and ORF7b help viral infection, trafficking, and maybe the budding of the virion; NSP1, NSP3, NSP5, NSP6, NSP7, and N protein suppress host cell replication, transcription, and translation, and at the same time contribute to the same processes in the viral lifecycle; S protein facilitates the formation of new viruses after viral replication and translation of structural proteins; NSP1, NSP5, N protein, M protein, and ORF7 may inhibit host cell antiviral responses and therefore promote the survival of the infected virus, and they also prolong host cell survival to increase viral production. NSP6 and NSP7 may manipulate host cell metabolism and signaling transduction pathways. Understanding in detail how the virus and host cell communicate will help us to find or design therapeutic strategies to suppress viral infection.

Additionally, we compared NSP1 and N protein interactomes among different human coronaviruses. These comparisons expand our knowledge of these viruses. Although NSP1 and N proteins from different coronaviruses share limited sequence identity, they have many common host binding partners, which match their biological functions. Such as the suppression of host gene expression by NSP1 proteins may be mediated through their binding to the EIF complex. The regulation of viral replication, transcription, and translation in all human coronaviruses may through the interaction between N protein and host ribosomal proteins or proteins involved in ribosomal biogenesis. Besides these common interaction proteins, we also uncovered some host proteins have different interaction ability among the seven human coronaviruses, and this may explain differing pathogenesis patterns among the coronaviruses. We identified and validated strong interactions between the N proteins of SARS-CoV-2 and SARS-CoV-1 with host G3BP1 or G3BP2, two proteins that are known to be involved in innate immunity. These strong interactions may indicate that N proteins in SARS-CoV-1/2 may suppress the host antiviral response and improve viral infection. Therefore, SARS-CoV-2 and SARS-CoV-1 may be more contagious than other coronaviruses.

In conclusion, our systemic study of the SARS-CoV-2 protein-protein interaction network provides useful data on viral gene/protein functions and potential underlying mechanisms, which could lead to the identification of new drug targets for the treatment of COVID-19. The comparison of interactomes among different human coronaviruses elucidated the common and different strategies these human coronaviruses may employ to manipulate host cells and maximize viral infection/production. These findings will benefit our current fight against COVID-19 and also suggest ways to combat any future coronavirus-caused diseases.

## MATERIALS and METHODS

### Plasmids, Cell Culture, and Transfection

The cDNAs of viral genes were synthesized using Integrated DNA Technologies or Twist Bioscience, or ordered from Addgene, and subcloned into pDONOR201 vector as entry clones. Then the entry clones were recombined into a lentiviral-gateway–compatible destination vector to express the fusion proteins with SFB or BioID2 tag.

HEK293T and U2OS cell lines were purchased from American Type Culture Collection. They were maintained in DMEM supplemented with 10% fetal bovine serum, and cultured at 37 °C in 5% CO_2_ (v/v).

Recombined destination vector with SFB- or BioID2-fused viral genes were transfected into HEK293T or U2OS cells with polyethyleneimine reagent (Fisher Scientific). After selection with 2 μg/ml puromycin (Life Technologies), the stable clones were picked and validated by Western blot analysis. The exception is the viral gene NSP1: destination vectors with NSP1 (SFB-NSP1 and BioID2-NSP1) were transfected into HEK293T cells 36 hours before cell harvest using the reagent X-tremeGENE HP DNA Transfection Reagent (Sigma-Aldrich). Cells expressing the BioID2-tagged genes were treated with 50mM biotin for 18 hours and then harvested. Vector-transfected cells or cells expressing GFP were used as controls, which were processed along with these experiments. Three biological replicates were performed for each SFB- or BioID2-tagged gene or negative control.

### SFB-TAP and BioID2 Assays

Pellets of cells expressing SFB- or BioID2-tagged viral genes were subjected to lysis with NETN buffer (100mM NaCl, 1mM EDTA, 20mM Tris-HCl, and 0.5% Nonidet P-40) with protease inhibitor cocktail (Sigma-Aldrich) at 4 °C for 20 minutes. After centrifugation at 14,000 rpm for 20 minutes at 4 °C, the supernatant was incubated with streptavidin-conjugated beads (Fisher Scientific) for 2 hours at 4 °C. Then the beads were washed with NETN buffer four times. The BioID2-tagged samples were subjected to SDS-PAGE, and the gel was fixed and stained with Coomassie brilliant blue. Then the whole lane of the sample in the gel was excised and subjected to MS analysis. The SFB-tagged samples were eluted by 2 mg/ml biotin for 1 hour at 4 °C. The elutes were incubated with S-protein agarose (VWR International) for 2 hours. After four washings with NETN buffer, the beads were subjected to SDS-PAGE and Coomassie brilliant blue staining, as with the BioID2 samples.

### Mass Spectrometry Analysis

The samples in gel were excised and de-stained completely before digestion. In-gel digestion was performed with trypsin (V5280, Promega Corporation) in 50mM NH_4_HCO_3_ at 37 °C overnight. The extracted peptides were vacuum-dried and then reconstituted in the MS loading solution (2% acetonitrile and 0.1% formic acid).

The MS sample was loaded onto nano-reverse-phase high-performance liquid chromatography and eluted with acetonitrile gradient from 5% to 35% for 60 minutes at a flow rate of 300 nL/minute. The elute was analyzed by the Q Exactive HF MS system (Thermo Fisher Scientific) with positive ion mode and in a data-dependent manner, one full scan followed by up to 20 MS/MS scans. The full MS scan was performed with a scanning range of 350-1200 m/z and resolution at 60,000 at m/z 400.

The raw MS data were submitted to Mascot 2.5 (Matrix Science) using Proteome Discoverer 2.2 (Thermo Fisher Scientific). The database used for searching was Homo sapiens downloaded from Uniprot (July 2020). The sequences of SARS-CoV-2 genes, SFB tag, and BioID2 tag were added into the database with 20,414 entries in total. Oxidation for methionine and carboxyamidomethyl for cysteine were set as variable modifications. The mass tolerance for precursor was 10 ppm, and for product ion, 0.02 Da. Tolerance of two missed cleavages of trypsin was applied. Common contaminant proteins were removed from the identification list. The mass spectrometry proteomics data have been deposited to the ProteomeXchange Consortium via the PRIDE partner repository with the dataset identifier PXD023209.

### High-Confidence Interacting Proteins

The identified protein lists were applied to a filtration strategy by comparing with lists obtained from controls to uncover HCIPs. We used SAINTexpress (version 3.6.3) to compare samples with the controls, which included the results from SFB-TAP or BioID2 experiments using the bait, vector, GFP control, and purification results with other virus genes, except the one we were analyzing. For the SARS-CoV-2 HCIPs analysis, we selected the proteins with SAINTexpress Bayesian false-discovery rate ≤0.05 as our virus interactome HCIPs. The host-virus interactome network was generated by Cytoscape and based on these HCIPs. Functional characterization was carried out by Metascape^65^ or Ingenuity Pathway Analysis (Qiagen).

### Pulldown and Western Blot Analysis

To validate the interaction between host and viral gene products, we transfected HEK293T cells with constructs encoding SFB-tagged viral genes using polyethyleneimine reagent. Cells were collected and subjected to lysis buffer (NETN buffer) on ice for 20 minutes. The cell lysates were collected and centrifuged. Supernatants were incubated with S-protein beads for 2 hours at 4 °C. After three washings with NETN buffer, samples were boiled in 2× Laemmli buffer and the Western blot analysis was conducted with antibodies as indicated in the figures.

### Immunofluorescent Staining

Immunofluorescent staining was performed following the method described in our previous study^74^. U2OS cells were cultured on coverslips overnight and then transfected with construct encoding SARS-CoV-2 viral gene or vector control. Twenty hours after transfection, cells were fixed in 3% paraformaldehyde for 10 minutes and extracted with a 0.5% Triton X-100 solution for 5 minutes. After blocking with 1% bovine serum albumin, cells were incubated with the indicated primary antibodies for 1 hour and washed three times before incubation with fluorescein isothiocyanate- or rhodamine-conjugated second primary antibodies (1:3000 dilution; Jackson ImmunoResearch Laboratories) for 1 hour. The coverslips were mounted onto glass slides with ProLong gold antifade reagent with DAPI (Cell Signaling Technology) and visualized with a fluorescent microscope.

### Luciferase Reporter Assay

The luciferase reporter assay followed a previous study^75^. Viral proteins were overexpressed in HEK293T cells along with IFN-beta-luciferase or 5× ISRE-Luc reporter, and with the pRL-Luc with Renilla luciferase as the internal control. Cells were lysed with passive lysis buffer (Promega), and we performed the luciferase assays using a dual-luciferase assay kit (Promega) and quantified them with Monolight 3010 (Becton Dickinson).

### Statistical Analysis

Each experiment was repeated three times or more unless otherwise noted. Differences between groups were analyzed using the Student *t* test or two-way analysis of variance with the Tukey multiple comparisons test. P values less than 0.05 were considered statistically significant.

## ACKNOWLEDGMENTS

We thank Drs. Peihui Wang and Yuan Wang for their kind help. We thank Erica Goodoff and the Research Medical Library at the University of Texas MD Anderson Cancer Center for editing the manuscript. This work was supported in part by startup and internal funds from MD Anderson Cancer Center to J.C. J.C. also received support from the Pamela and Wayne Garrison Distinguished Chair in Cancer Research. This work was also supported in part by the National Institutes of Health through Cancer Center Support Grant P30CA016672. In addition, JC receives support from Cancer Prevention & Research Institute of Texas award (RP160667 and RP180813) and NIH (P01CA193124, R01CA210929, R01CA216437 and R01 CA216911).

## AUTOHOR CONTRIBUTIONS

Conceptualization, Z.C. and J.C.; Methodology, Z.C., C.W., X.F., and J.C.; Investigation, Z.C., C.W., and X.F.; Validation, Z.C., C.W., and X.F.; Resources, L.N., M.T., H.Z., Y.X., S.K.S., and M.S.; Writing Original Draft, Z.C. and J.C.; Review and Editing, all authors; Supervision, J.C.

## Supplemental Figure Legends

**Figure S1, related to Figure 1.**
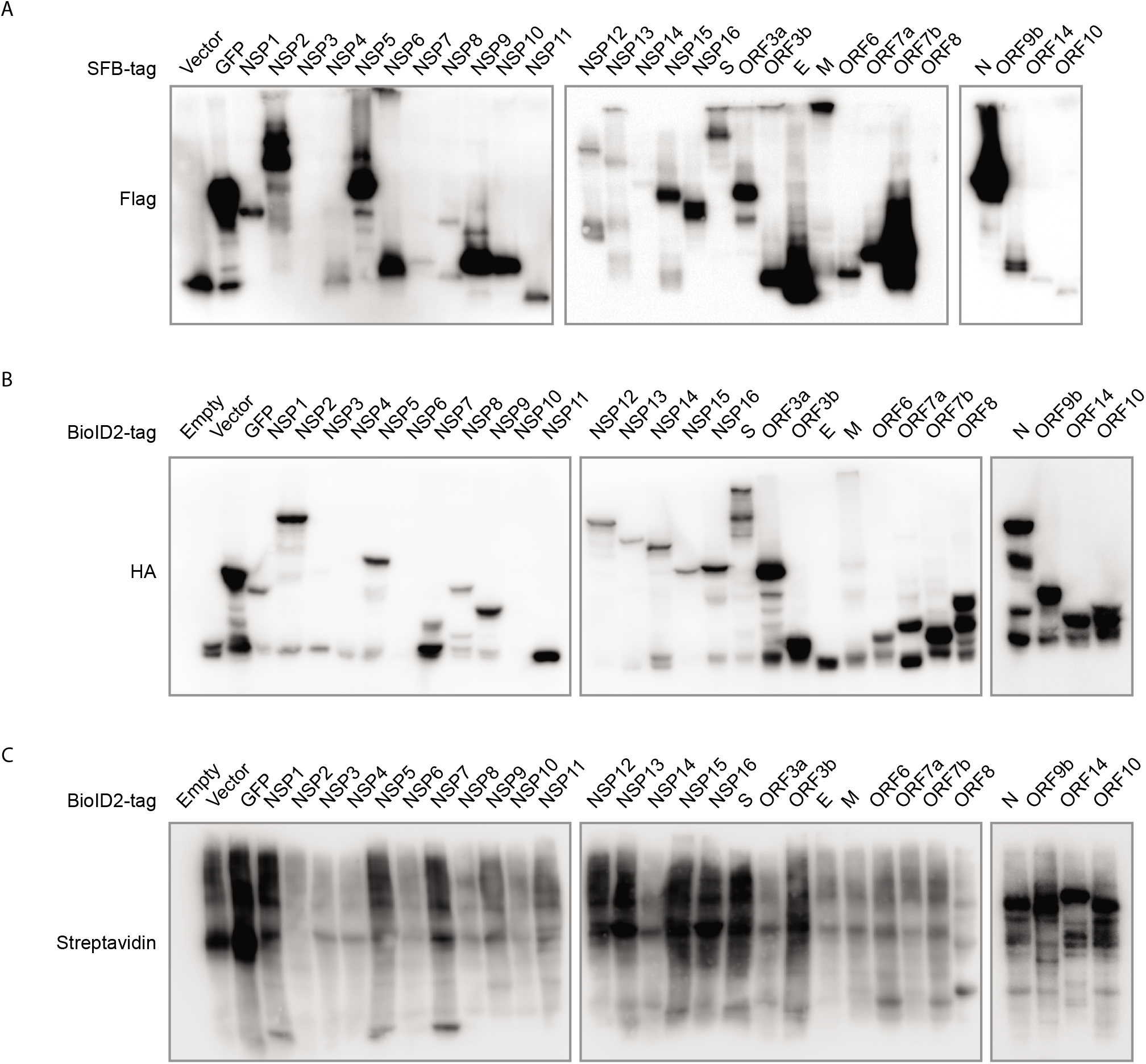
Validation of SARS-CoV-2 protein expression. (A) Twenty-nine SARS-CoV-2 genes with the SFB tag were expressed in cells. Cells transfected with vector or construct encoding SFB-tagged GFP were included as controls. (B) Twenty-nine SARS-CoV-2 genes with the second-generation biotin ligase (BioID2) tag were expressed in cells. Untransfected cells, cells transfected with vector, and cells transfected with construct encoding GFP tagged with BioID2 were included as controls. (C) Validation of the BioID2 labeling with staining of the streptavidin antibody.

**Figure S2, related to Figure 1.**
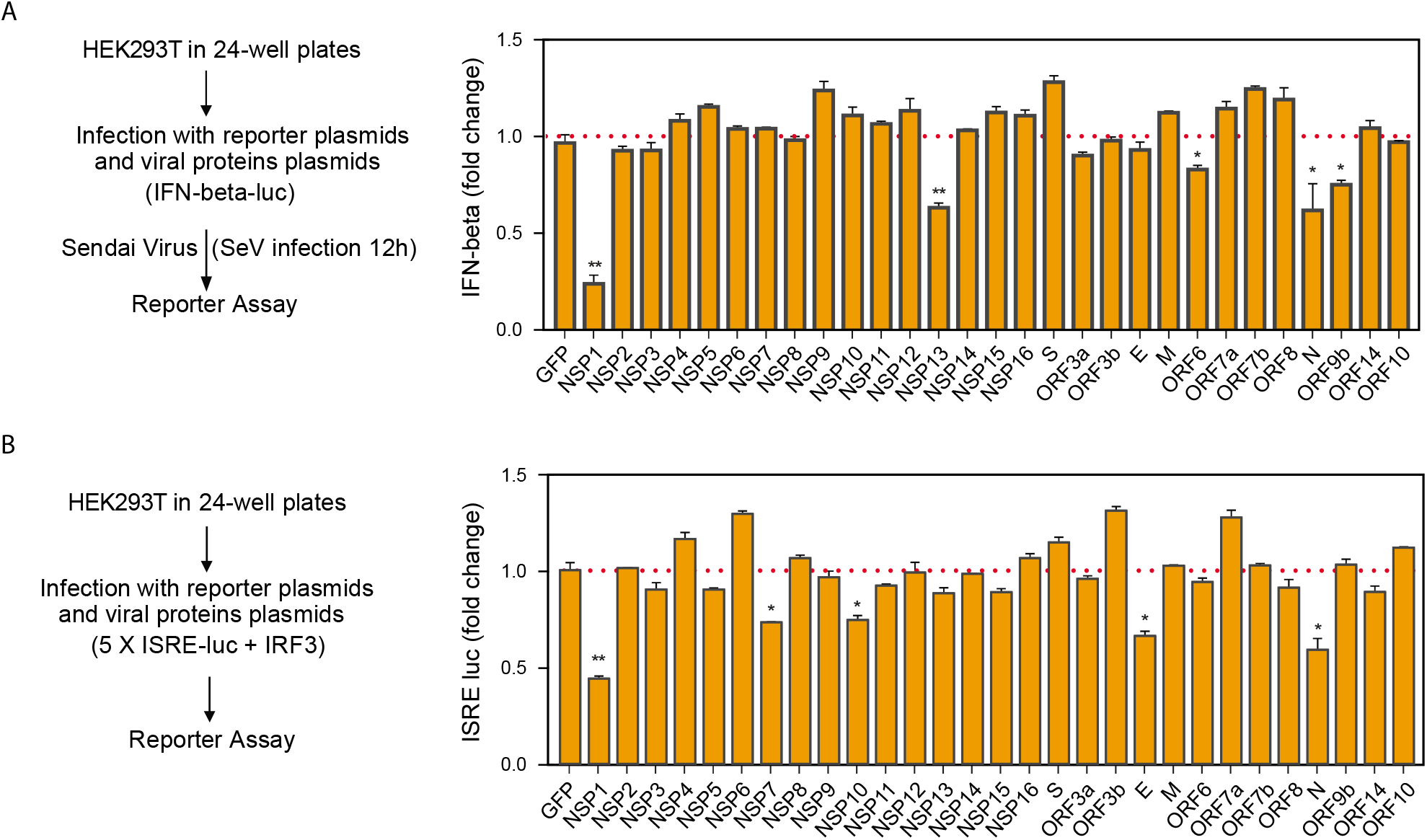
Interferon (IFN) signaling assay for SARS-CoV-2 coding genes. (A) Analysis of IFN-beta-luciferase reporter 12 hours after Sendai virus (SeV) infection. (B) Analysis of ISRE-luciferase reporter when the viral proteins were co-expressed with IRF3. Graphs show mean ± SD, n = 3. **p < 0.01 and *p < 0.05 (Student’s *t*-test).

**Figure S3, related to Figure 5.**
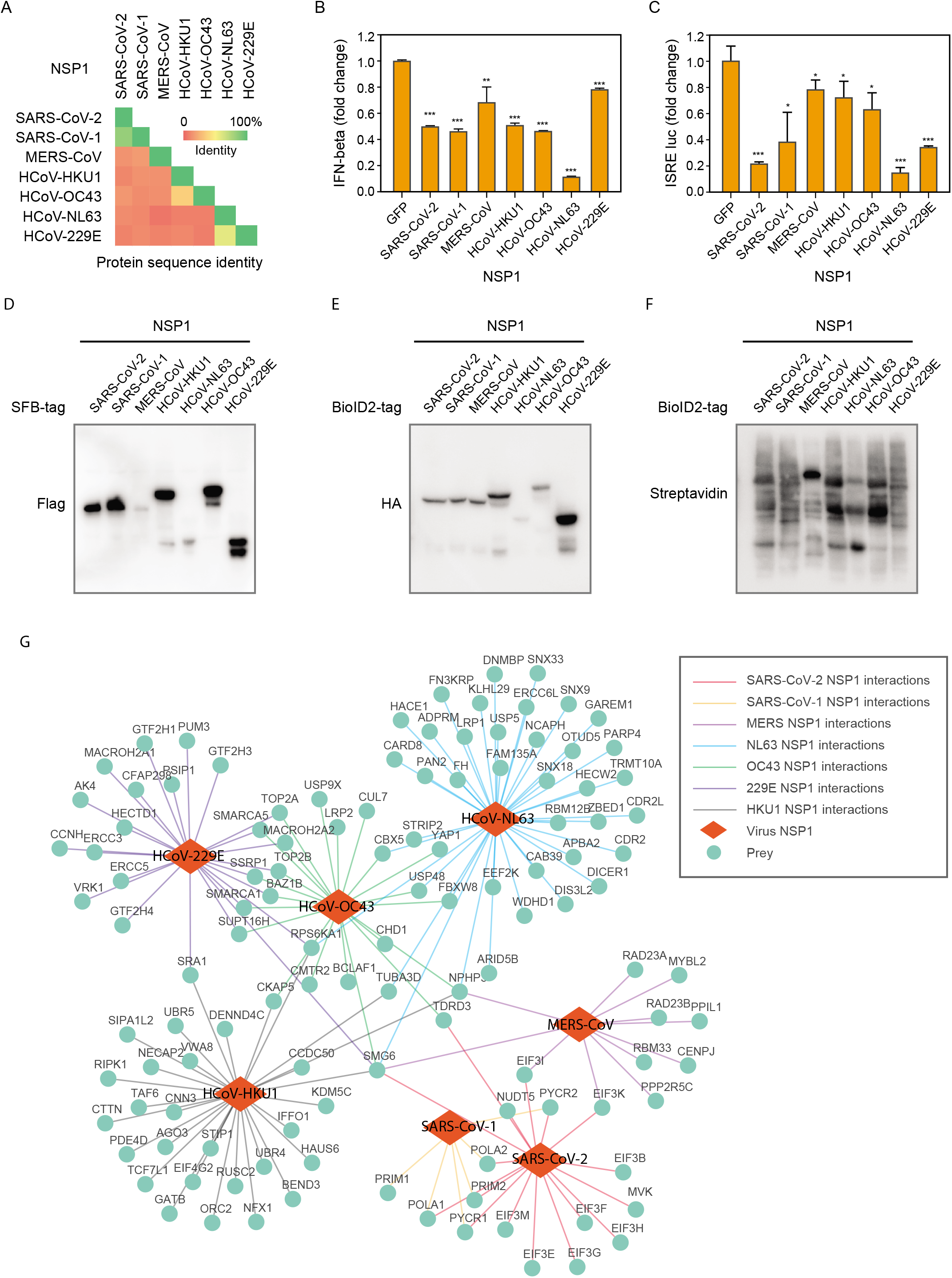
Comparison of interactomes of seven NSP1 proteins from seven human coronaviruses. (A) Sequence identity comparison of NSP1 proteins from seven human coronaviruses. (B, C) Interferon (IFN) signaling assay for NSP1 of seven human coronaviruses. (B) Ana lysis of IFN-beta-luciferase reporter 12 hours after Sendai virus infection. (C) Analysis of ISRE luciferase reporter when the viral proteins were co-expressed with IRF3. Graphs show mean ± SD, n = 3. ***p < 0.001, **p < 0.01 and *p < 0.05 (Student’s *t*-test). (D) Expression validation of NSP1 with the SFB tag from different human coronaviruses. (E) Expression validation of NSP1 with BioID2 tag from different human coronaviruses. (F) Validation of labeling efficiency for the BioID2-tagged gene products. (G) Network of high-confidence interactive proteins (green circles) identified with seven NSP1 proteins (red diamonds).

**Figure S4, related to Figure 6.**
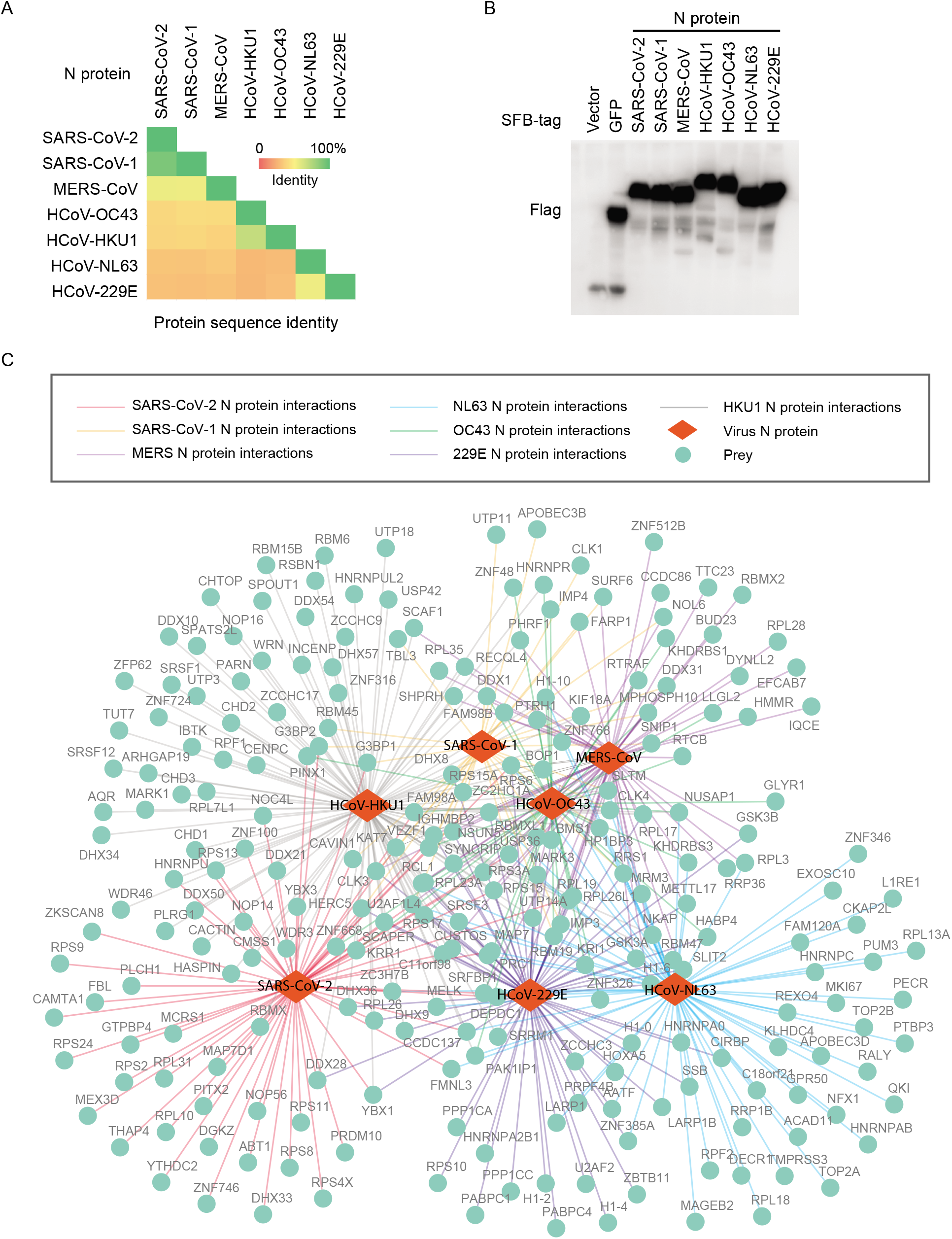
Comparison of interactomes of seven N proteins from seven human coronaviruses. (A) Sequence identity comparison of N proteins from seven human coronaviruses. (B) Expression validation of NSP1 with the SFB tag from different human coronaviruses. (C) Network of high-confidence interactive proteins (green circles) identified with seven NSP1 proteins (red diamonds).

**Figure S5.**
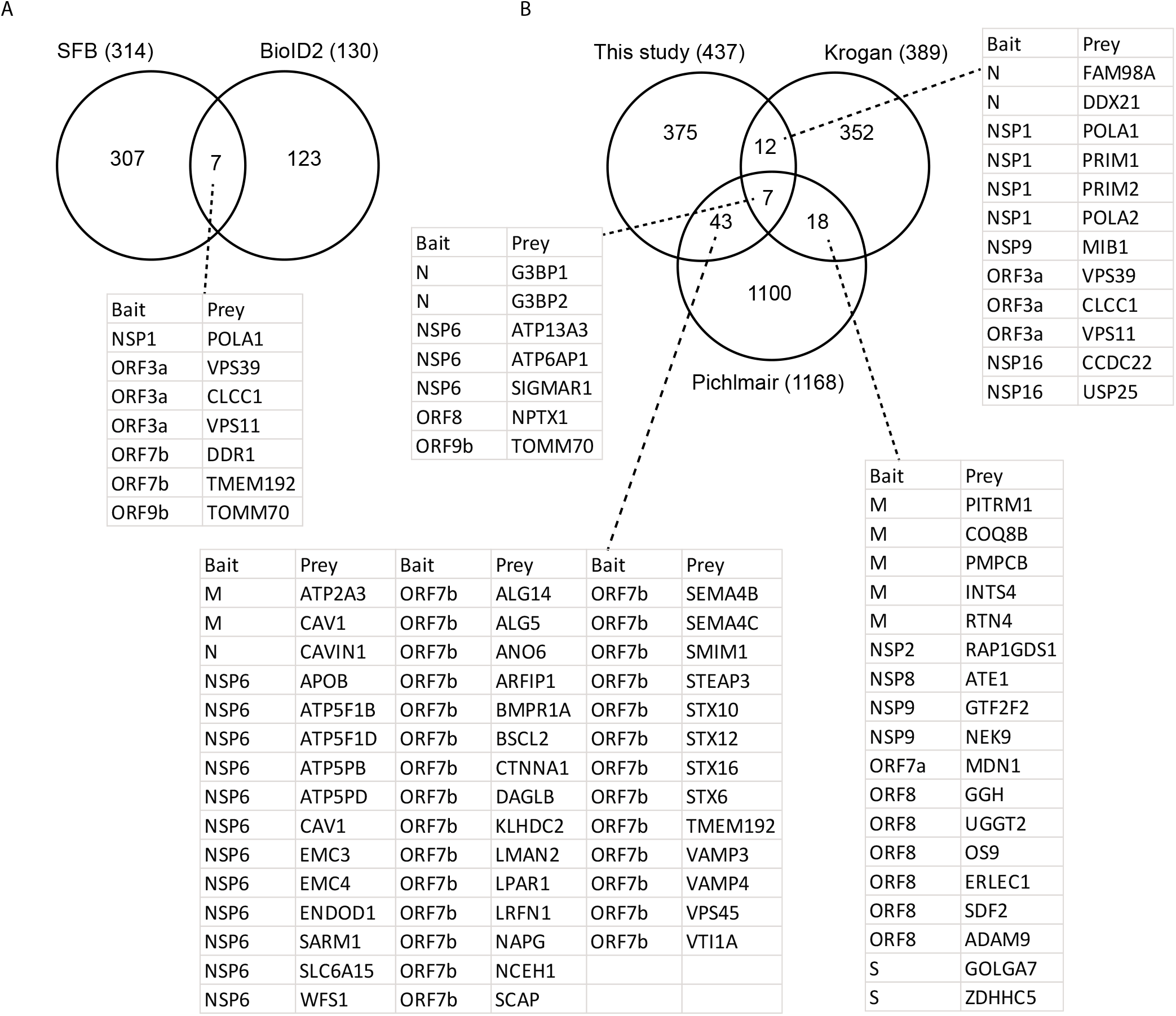
Overlap of high-confidence interacting proteins (HCIPs) from different datasets. (A) Overlap of HCIPs identified from SFB-tandem affinity purification and second-generation biotin ligase (BioID2) labeling experiments. (B) Overlap of HCIPs reported previously and those identified in the current study.

## Supplemental Table Legends

**Table S1, related to Figure 1. Raw identifications of all SFB-TAP and BioID2 purifications.**

**Table S2, related to Figure 1. HCIPs of SFB-TAP and BioID2 purifications.**

**Table S3, related to Figure 4. Drugs target human proteins which binding to SARS-CoV-2 viral proteins.**

**Table S4, related to Figure 5. NSP1 interactomes raw identifications and HCIPs of all seven human coronaviruses using SFB-TAP purification.**

**Table S5, related to Figure 5. NSP1 interactomes raw identifications and HCIPs of all seven human coronaviruses using BioID2 purification.**

**Table S6, related to Figures 6 and 7. N proteins interactomes raw identifications and HCIPs of all seven human coronaviruses using SFB-TAP purification.**

